# Homeostatic balance of Gut-resident Tregs (GTregs) plays a pivotal role in maintaining bone health under post-menopausal osteoporotic conditions

**DOI:** 10.1101/2024.09.13.612848

**Authors:** Asha Bhardwaj, Leena Sapra, Divya Madan, Vineet Ahuja, Pradyumna K. Mishra, Rupesh K. Srivastava

## Abstract

Osteoporosis is a skeletal condition characterized by the deterioration of bone tissue. The immune system plays a crucial role in maintaining bone homeostasis and combating the development of osteoporosis. Immunoporosis is the term used to describe the recent convergence of research on the immune system’s role in osteoporosis. Gut harbors the largest component of the immune system and there is growing evidence that intestinal immunity plays a vital role in regulating bone health. Gut-resident regulatory T cells (GTregs) play an essential role in inhibiting immune responses and preventing various inflammatory manifestations. Our findings show that GTregs have a pivotal role in the pathophysiology of post-menopausal osteoporosis (PMO). We investigated the potential of GTregs in regulating the development of bone cells *in vitro*. We observed that GTregs significantly enhance osteoblastogenesis with concomitant inhibition of osteoclastogenesis in a cell-ratio-dependent manner. We further report that the deficiency of short-chain fatty acids (SCFAs) in osteoporotic conditions substantially disrupts the composition of GTregs, leading to a loss of peripherally derived Tregs (pTregs) and an expansion of thymus-derived Tregs (tTregs). Moreover, the administration of probiotics *Lactobacillus rhamnosus* and *Bifidobacterium longum* modulated the GTregs compartment in an SCFA-dependent manner to mitigate inflammatory bone loss in PMO. Notably, SCFAs-primed GTregs were found to be significantly more effective in inhibiting osteoclastogenesis compared to unprimed GTregs. Altogether our results, for the first time, highlight the crucial role of GTregs in the pathophysiology of PMO, with potential clinical implications in the near future.

**Graphical Abstract:**
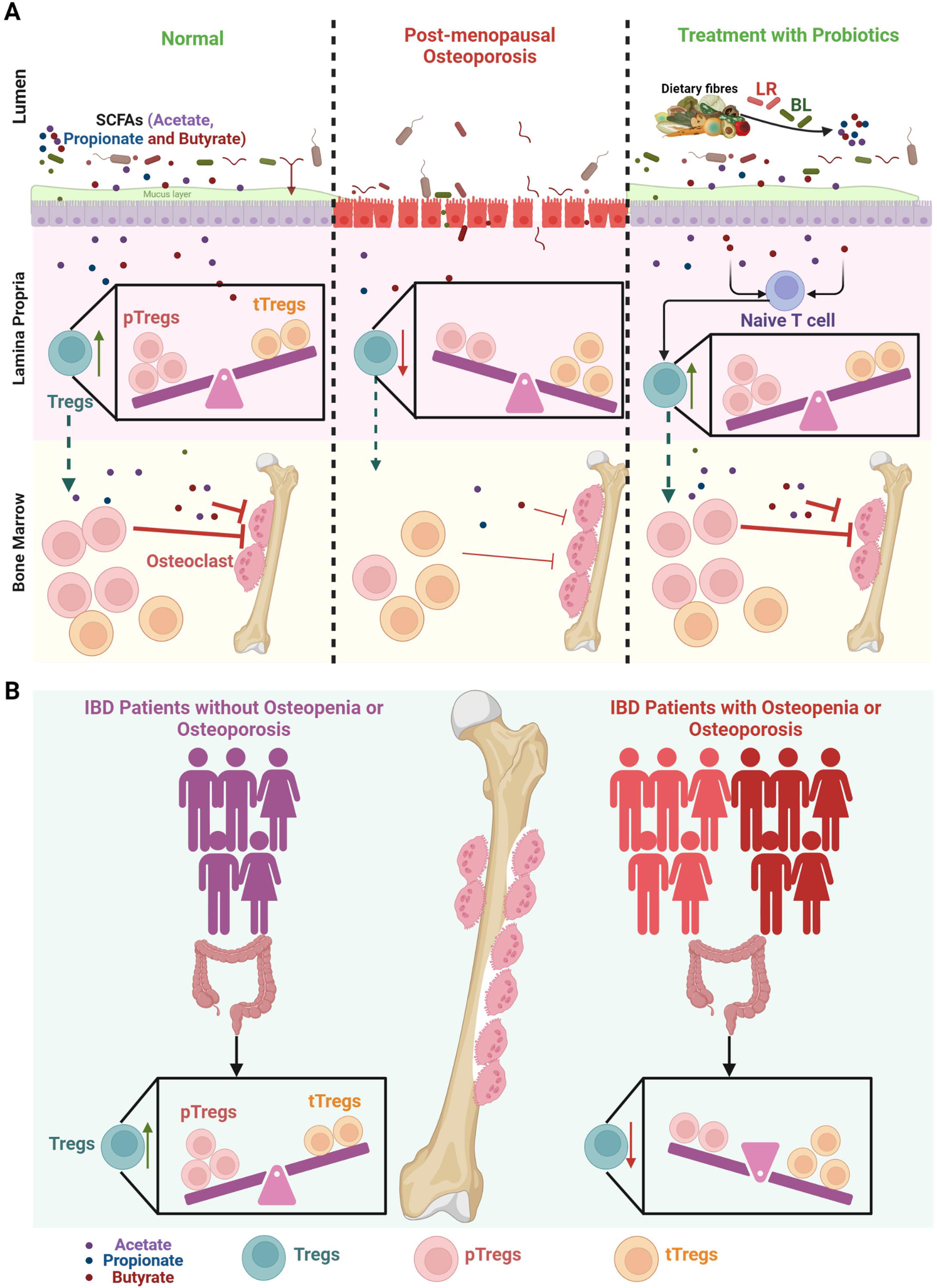
A) During normal physiological conditions there is a sufficient proportion of Tregs in the mice intestine with a higher frequency of pTregs compared to the tTregs. Gut-resident Tregs inhibit osteoclastogenesis and therefore prevent bone loss. However post-menopausal condition decreases the Treg population in the intestinal tissues and further perturbs the balance of pTregs and tTregs resulting in bone loss. Probiotics such as *Lactobacillus rhamnosus* (LR) and *Bifidobacterium longum* (BL) induce the development of Tregs from naïve T cells and further restore the balance of pTregs and tTregs in a short chain fatty acid (SCFA) dependent manner. SCFAs primed Tregs are more potent in inhibiting osteoclastogenesis thereby preventing bone loss due to osteoporosis. B) Similar to mice the frequency of Tregs decreases in the human colon of inflammatory bowel disease (IBD) patients with osteopenia or osteoporosis compared to the IBD patients without osteopenia/osteoporosis

## 1.0 Introduction

Osteoporosis is an inflammatory skeletal disease characterized by the deterioration of bone tissues, reduction in bone strength, bone microarchitecture, and bone mineralization with a huge socio-economic burden ^1,2^. After ischemic heart disease, dementia, and lung cancer, it is the fourth most severe chronic condition and an increasing public health concern (IOF 2021). The immune system plays a crucial role in the development of osteoporosis. In lieu of the growing involvement of the immune system in the pathophysiology of osteoporosis, our group has coined the term “Immunoporosis”^2–5^. We reported that the pathophysiology of osteoporosis is largely mediated via regulatory T cells (Treg) ^6–10^, as they exhibit the potential to suppress osteoclastogenesis and thus enhance bone formation ^11–13^.

Fascinatingly, it has been observed that the gut harbors 70-80 % of the immune system in the human body ^14–16^. Moreover, Tregs (approximately 30% of the total CD4 population) primarily reside in the intestine (GTregs), and any aberration in the population of GTregs or their associated cytokines is related to several diseases including bone pathologies such as spondyloarthritis ^17,18^. Nevertheless, the involvement of GTregs in post-menopausal osteoporosis (PMO) has not been investigated to date. Thus, in the present study, we investigated the role of GTregs in PMO.

In a healthy gut, Tregs play a crucial role in maintaining intestinal homeostasis by inhibiting unwarranted immune responses ^19^. Through a variety of molecular processes, GTregs regulate the mucosal immune responses at several cellular levels. Various evidence points to the existence of two developmentally distinct subsets of Tregs, thymus-derived Tregs (tTregs) and peripherally derived Tregs (pTregs), which develop as naïve CD4^+^ T cells in the thymus but become Tregs in the peripheral tissues in response to gut microbiota and their metabolites ^17^. Unlike pTregs, tissue-resident tTregs constitutively express helios and the cell surface marker neuropilin 1 (NRP-1) ^17^ and thus NRP-1 and helios serve as valuable markers for identifying tTregs and distinguishing them from pTregs ^17^. tTregs are vital to preserving immunological homeostasis and tolerance to the self whereas pTregs regulate the immunity at sites of inflammation ^20^. The milieu in the gut favors the development of pTregs ^17^. Recent advances have reported that PMO is associated with dysbiosis and increased gut permeability ^21–23^. We therefore hypothesized that gut dysbiosis and increased gut permeability during osteoporosis would drastically affect the intestinal Treg cell homeostasis. Moreover, various gut-associated bacterial metabolites are important in modulating this homeostasis.

Surprisingly, several pieces of evidence suggest that *Lactobacillus rhamnosus* (LR) and *Bifidobacterium longum* (BL) via preserving the intestinal integrity and preventing gut dysbiosis enhance the anti-osteoclastogenic subset of Tregs ^10,24–26^. Nevertheless, the potential of these probiotics in regulating the homeostasis between pTregs and tTregs remains unexplored. Thus, in this study, we further investigated the role of probiotics LR and BL in alleviating inflammatory bone loss via their effect on GTreg homeostasis.

In the present study, we for the first time report that a deficiency of short-chain fatty acids (SCFAs) under ovariectomized conditions significantly decreases both the number and the composition of GTregs (reduced pTregs and enhanced tTregs). GTregs promote osteoblast formation along with significantly suppressing osteoclast formation. Interestingly, the probiotics LR and BL restored the GTregs homeostasis by inducing the differentiation of pTregs in a SCFAs-dependent manner thereby preventing bone resorption. Overall, our study elucidates the cellular mechanisms that are vital in shaping the Tregs landscape in the gut under PMO conditions.

## 2.0 Methods

### 2.1 Reagents and Antibodies

The following antibodies and kits were procured from eBioscience (San Diego, CA, USA): PerCp-Cy5.5 Anti-Mouse-CD4-(RM4-5) (550954), APC Anti-Mouse/Rat-Foxp3 (FJK-16s) (17-5773), PEcy7 Anti-Mouse-NRP-1 (3DS304M) (25-3041), Foxp3/Transcription factor staining buffer (0-5523-00), Anti-IFN-γ (clone: XMG1.2), Anti-IL-4 (clone: 11B11), and RBC lysis buffer (00-4300-54). The PEcy7 Anti-Human-Helios (22F6) (137235), FITC Anti-Human-CD3 (UCHT1)(300406), PE-Anti-Human-CD4 (RPA-T4)(300550), and Alexa Flour 647 Anti-Human-Foxp3 (249D) (320214), were procured from BioLegend. The acid phosphatase leukocyte (TRAP) kit (387A), ascorbic acid (A4544), and β-glycerophosphate (G9422) were purchased from Sigma (St. Louis, MO, USA). Macrophage-colony stimulating factor (M-CSF) (300-25) and receptor activator of nuclear factor κB-ligand (RANKL) (310-01), human TGF-β1 (AF-100-21C) and murine IL-2 (AF-212-12) (200-23) were procured from PeproTech (Rocky Hill, NJ, USA). α-Minimal essential media (MEM) and Roswell Park Memorial Institute (RPMI)-1640 media were purchased from Gibco (Grand Island, NY, USA). LR UBLR-58 and BL UBBL-64 were procured from Unique Biotech Ltd., Hyderabad, India.

### 2.2 Animals

Female C57BL/6J mice aged 8–10 weeks were used for both *in vitro* and *in vivo* investigations. All mice were kept in the animal facility of All India Institute of Medical Sciences (AIIMS), New Delhi, India under specific pathogen-free conditions and provided continuously with sterilized food and autoclaved drinking water ad libitum. Mice were either sham-operated or ovariectomized (ovx) after anesthetizing with ketamine (100–150 mg/kg) and xylazine (5–16 mg/kg). Afterward, mice were randomly assigned to either two experimental groups’ viz. sham and ovx or four groups viz. sham, ovx, ovx + LR, and ovx + BL. Mice in the ovx + LR and ovx + BL groups were gavaged 200 µl (10^9^ CFU) of LR and BL suspension respectively daily for 45 days starting one week after surgery. At the end of the experiment, mice were euthanized and several tissues such as blood, bone, mesenteric lymph node (MLN), and intestine were harvested. All medical procedures were performed on mice after due approval from the Institutional Animal Ethics Committee of AIIMS, New Delhi, India (85/IEAC-1/2018).

### 2.3 Human subjects

20 subjects (14 males and 6 females with an average mean age of 31 years) with confirmed inflammatory bowel disease (IBD) either ulcerative colitis (16) or Crohn’s disease (4) undergoing treatment in AIIMS, New Delhi were recruited in the study (AIIMSA1342). Dual-energy X-ray absorptiometry (DEXA) scans were performed on the patients and were categorized into three groups: Control IBD patients (without osteopenia and osteoporosis) with a T score of more than −1.0, IBD patients with osteopenia (T score between −1.0 to −2.5) and IBD patients with osteoporosis (T score of less than −2.5). Biopsies of the sigmoid colon were assessed in all groups and processed for flow cytometry.

### 2.4 Analysis of short-chain fatty acids (SCFAs)

SCFA content in the fecal matter of mice was determined with the help of high-performance liquid chromatography (HPLC). For the same 300 mg of the fecal sample was weighed and 1 ml of Milli Q and 100 µl of hydrochloric (HCl) acid were added to the fecal sample. The fecal sample was then completely homogenized with the help of the vortex (2-3 minutes) and placed for 20 minutes with shaking of the sample twice in 20 minutes. After 20 minutes samples were centrifuged at 13850 g for 10 minutes at 4° C. Supernatant was collected and transferred to the 2 ml Eppendorf. 600 µl of the diethyl ether was added to the supernatant and continued extraction for 20 minutes. After 20 minutes of extraction centrifuged the sample at 850 g for 5 minutes at 4° C. Collected 400 µl of the supernatant or organic layer and added to the 500 µl of the sodium hydroxide (NaOH). Continued extraction for 20 minutes and subsequently centrifuged the sample at 850 g for 5 minutes. Discarded the ether layer or organic layer and collected 450 µl of the aqueous layer or water-soluble layer. Then added water-soluble layer to the 300 µl of HCl and immediately filtered the sample through the 0.22 µm filter. Samples were run on an HPLC machine. HPLC conditions used were: Xterra C18 column (250 mm X 4.6 mm X 3.5 µm); mobile phase, 10 mM sulphuric acid (H_2_SO_4_) (Isocratic gradient); flow rate, 0.6 ml/min; run time: 11 minutes, wavelength: 210 nm. SCFA level was determined using the standard calibration curves.

### 2.5 Micro-Computed Tomography (µ-CT) measurements

As previously mentioned, an *in vivo* X-ray SkyScan 1076 scanner (Aartselaar, Belgium) was used to perform µ-CT scanning and analysis ^6–10,26,27^. Briefly, samples were oriented correctly in the sample holder and scanned at 50 kV, 204 mA, with a 0.5-mm aluminum filter. Software called NRecon was used for the reconstruction procedure. Following reconstruction, ROI was created in secondary spongiosa at 1.5 mm from the distal edge of growth plates using a total of 100 slices. This ROI was then processed using CTAn application for determining the several micro-architectural parameters of bone samples. The bone mineral density (BMD) of the LV-5 trabecular, femur trabecular, tibia trabecular, femur cortical, and tibia cortical region was calculated using the volume of interest of µ-CT scans performed on the trabecular regions. BMD was assessed using calibrators made of 4-mm-diameter hydroxyapatite phantom rods with known BMD (0.25 g/cm^3^ and 0.75 g/cm^3^) ^27^.

### 2.6 Immune cells isolation from mice intestinal tissues

After isolating MLN, small intestine (SI), and large intestine (LI) were collected. MLN was minced with the help of frosted end slides and passed through the 70 µM cell strainer. The resulting cell suspension was then used for cell staining. For isolating immune cells from the SI, Peyer’s patches were removed and then cells were isolated as per the protocol of lymphocyte isolation from the SI by Qiu et al. ^28^. Briefly, small intestinal tissues were shaken at 220 rpm for 20 minutes at 37 °C in Hank’s balanced salt solution (HBSS) (TL1109) containing 1mM dithioerythritol (GRM359). Repeated the above step twice to remove the intraepithelial lymphocytes. The remaining tissues were further processed with shaking at 220 rpm for 30 minutes at 37 °C in HBSS containing 1.3 mM ethylenediaminetetraacetic acid (EDTA) (12135). The step was repeated twice to remove the epithelial cells. Tissues were then washed and digested with RPMI containing collagenase type 1 (100 U/ml) (17100017) by shaking at 220 rpm for 45 min at 37 °C for isolation of lamina propria (LP) cells. After digestion, lymphocytes were harvested using a 44% and 67% Percol gradient purification. For the isolation of lymphocytes from the LI caecum was removed and the colon was processed similarly to the SI.

### 2.7 Immune cell isolation from human colon tissue

Colonic biopsy was treated twice with 1mM of EDTA solution in 1 X HBSS at 37°C and 220 rpm for 30 min to remove the epithelial cells. The remaining tissue was washed twice with 1XHBSS for 15 minutes and digested with 2mg/ml collagenase type 1 solution in RPMI media for 1-2 hours at 37°C and 220 rpm. The digested tissue was passed through the 70 µm filter and washed with RPMI media.

### 2.8 Flow cytometry

Cells were harvested from the bone marrow (BM), MLN, and intestine (both SI and LI) of mice and stained for Tregs. For surface staining, cells were initially labeled with anti-CD4-PerCPcy5.5 and anti-NRP-1-PECy7 and incubated in the dark for 30 minutes in ice. Cells were washed and then fixed and permeabilized using the 1X-fixation-permeabilization buffer in the dark for 30 minutes in ice before intracellular staining. After washing cells were stained for anti-FoxP3-APC for 45 minutes. Lastly, cells were acquired on BD LSR fortessa (USA) and data was analyzed with the help of Flowjo-10 software (Treestar, USA). For staining of human Tregs, cells were first surface stained with anti-CD3-FITC, and anti-CD4-PE and incubated in the dark for 30 minutes in ice. Cells were then fixed, permeabilized, and stained for anti-FoxP3-APC and anti-Helios-PECy7 for 45 minutes. Cells were then acquired on BD Symphony and data was analyzed with the help of Flowjo-10 software. The gating strategy was adopted as per the experimental requirements.

### 2.9 Osteoclast differentiation and TRAP staining

C57BL/6J mouse aged 8 to 12 weeks was used to obtain bone marrow cells (BMCs) for osteoclast development by flushing the femoral bone with complete α-MEM media (10% heat-inactivated fetal bovine serum-FBS). BMCs were grown overnight in a T-25 flask in endotoxin-free complete α-MEM media supplemented with MCSF at the 35-ng/ml concentration after conducting red blood cell (RBC) lysis with 1XRBC lysis buffer. The following day, non-adherent cells were collected and co-cultured in a 96-well plate with 50,000 cells per well in the presence and absence of 0.3 mM conc. of SCFAs (acetate, propionate, and butyrate) in complete α-MEM supplemented with MCSF (30 ng/ml) and RANKL (60 ng/ml) for 4 days. On day 3 half of the media was replenished with the complete α-MEM media containing the factors. Lastly, tartrate acid-resistant phosphatase (TRAP) staining was performed as per the manufacturer’s instructions to examine the development of multinucleated osteoclasts. Briefly, at the end cells were carefully washed thrice with 1X phosphate-buffered saline (PBS), fixed, and then incubated for 10 minutes. After that, fixed cells were rinsed twice with 1XPBS and stained for 5–15 minutes in the dark at 37°C using a TRAP-staining solution. An inverted microscope (EVOS, Thermo Scientific, Waltham, MA, USA) was used to count and image multinucleated TRAP-positive cells with more than 3 nuclei, which were classified as osteoclasts. ImageJ software was used to estimate the osteoclast area and number (NIH, USA).

### 2.10 *In-vitro* differentiation of GTregs

Naïve CD4^+^ T cells were isolated from the MLN of 8-week-old C57BL/6J mice by using the T cell enrichment cocktail (BD). Negatively selectively cells were then seeded in 48 well plates coated with anti-CD3 (10 µg /ml) and anti-CD28 (2 µg/ml) and further stimulated with anti-IL-4 (5 µg/ml), anti-IFNγ (5 µg/ml), TGF-β1 (5 ng/ml) and IL-2 (10 ng/ml) in the presence and absence of 0.3 mM concentration of SCFAs (acetate, propionate, and butyrate) for 4 days. On day 5, cells were harvested and flow cytometry was performed to analyze the CD4^+^Foxp3^+^NRP^+^ and CD4^+^Foxp3^+^NRP^−^ cells.

### 2.11 Coculture of GTregs with BMCs for Osteoclastogenesis

For estimating the potential of GTregs in inhibiting osteoclastogenesis BMCs were cocultured with either GTregs or SCFAs-primed GTregs in different ratios (1:10, 1:5, and 1:1) in complete α-MEM media supplemented with MCSF (30 ng/ml) and RANKL (60 ng/ml) for 4 days. On day 3 half of the media was replenished with the complete α-MEM media containing the factors. Lastly, TRAP staining was performed to examine the development of multinucleated osteoclasts.

### 2.12 Coculture of Tregs with BMCs for Osteoblastogenesis

C57BL/6J mouse aged 8 to 12 weeks was used to obtain BMCs for osteoclast development by flushing the femoral bone with complete α-MEM media (10% heat-inactivated FBS). 50,000 BMCs per well were seeded in the 96 well plate in the presence of osteogenic induction media (OIM) consisting of the α-MEM media supplemented with β-glycerophosphate (10 mM) and ascorbic acid (50 µg/ml) for 24 hours. The following day half of the media was replenished with fresh OIM. On the 3^rd^ day 80% of the media was replenished with the fresh OIM. The next day, complete media was replenished and the BMCs were cocultured with the Tregs or SCFAs-primed Tregs in different ratios (1:10, 1:5, and 1:1) in the presence of OIM. On the 7^th^ day, alkaline phosphatase (ALP) staining was performed to examine the osteoblastogenesis with media replenishment every 3^rd^ day. For ALP staining culture media was removed from the plates and cells were washed two times with 1XPBS. 100 µl of the 10% formaldehyde was added to each well for 30 minutes at room temperature to fix the cells. After fixation cells were again washed with 1XPBS twice. 50 µl nitro blue tetrazolium (NBT)/ 5-bromo-4-chloro-3-indolyl-phosphate (BCIP) substrate was then added to each well and the plate was incubated at 37° C until the stain developed.

### 2.13 Gut permeability assay (Evans Blue Assay)

Evans Blue dye (30mg/kg of mice weight) was injected intravenously into the tail vein of mice. After half an hour of administration mice were sacrificed and the LI was harvested and kept in formamide solution at 72° C for 24 hours. Further, the supernatant was collected and absorbance was measured at 620 nm.

### 2.14 Histology

For histological analysis, SI, and LI were harvested from both sham and ovx groups. Tissues were fixed in 10% formaldehyde solution at room temperature, overnight for histological staining. After paraffin sectioning of tissues, slides were stained with hematoxylin and eosin. Slides were hydrated after the complete removal of wax which was followed by nuclear staining (hematoxylin). The excess stain was removed by rinsing the slide under tap water. Counterstaining via Eosin was performed which was next followed by dehydration and mounting. Tissue histological changes were identified using a microscope equipped with a digital camera.

### 2.15 Statistical Analysis

Statistical differences between groups (mice) were assessed using analysis of variance (ANOVA), followed by a paired or unpaired Student’s t-test as suitable. All of the data values are expressed as mean ± SEM. Differences among groups for human data were evaluated using two-tailed Mann–Whitney nonparametric and Student’s t-test for unpaired samples. Statistical significance was defined as p≤ 0.05 (*p<0.05, **p<0.01, ***p<0.001) to the groups indicated.

## 3.0 Results

### 3.1 GTregs are critically dysregulated during PMO

To investigate the possible involvement of GTregs in PMO, female C57BL6/J mice were randomly divided into two groups: sham (control) and ovx (ovariectomized) **(Figure 1A)**. On day 45 mice were sacrificed and bones were collected for µ-CT analysis. µ-CT results with significant changes in the bone histomorphometric parameters demonstrated that ovariectomy considerably deteriorates bone micro-architecture of the trabecular (LV-5, femur, and tibia) and cortical (femur and tibia) region in the ovx group in contrast to the sham group **(Figure 1B-C)**. Likewise, a significantly lower BMD at these sites in the ovx group compared to the sham group further supported the successful development of the PMO mice model **(Figure 1D)**.

**Figure 1:**
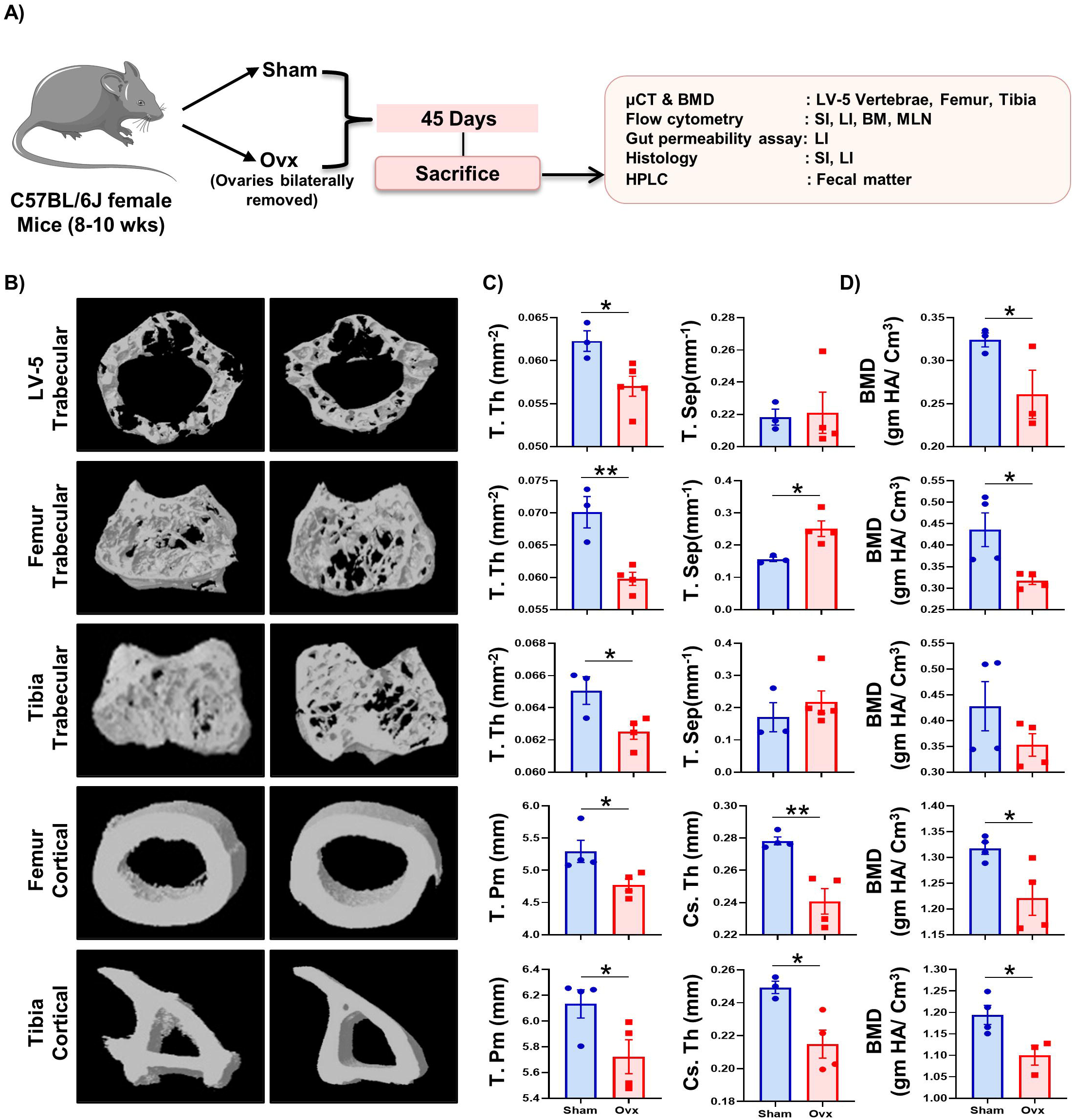
Development of post-menopausal osteoporotic mice model. A) Mice were divided into 2 groups: sham (control) and ovx (ovariectomized). At the end of 45 days, mice were sacrificed and analyzed for various parameters. B) 3D µ-CT reconstruction, C) Histomorphometric indices, and D) Graphical representation of bone mineral density (BMD) of LV-5 trabecular, femur trabecular, tibia trabecular, femur cortical, and tibia cortical regions of the sham and ovx groups. Trabecular thickness (Tb. Th); trabecular separation (Tb. Sep.); total cross-sectional perimeter (T. Pm); cross-sectional thickness (Cs. Th). Statistical significance was defined as *p ≤ 0.05, **p < 0.01 ***p ≤ 0.001 for the indicated mice group.

Next to determine the effect of PMO on intestinal health, firstly we assessed the length of the colon in both groups. We observed no discernible differences between the colon lengths of the two groups **(Figure 2A)**. However, histology data showed alterations in both the SI and LI, with more deterioration of villi structures in the ovx mice in comparison to the sham group **(Figure 2B**). Subsequently, compared to the sham, the LI of the ovx mice showed significantly enhanced intestinal permeability (p<0.05) **(Figure 2C)**. Further, to determine the status of GTregs in PMO, immune cells were harvested from the LP (prime site of Tregs) of both small intestine (LP-SI) and large intestine (LP-LI) and analyzed for Tregs (CD4^+^Foxp3^+^) **(Supplementary Figure 1).** In addition to LP-SI and LP-LI, MLN and BM (prime site of osteoclastogenesis) were also examined for Treg cells. Remarkably, in both the LP-SI and LP-LI, the frequency of GTregs was significantly lower (p<0.05 & p<0.01) in the ovx group as compared to the control group **(Figure 2D-E)**. Furthermore, the percentage of GTregs in the SI and LI showed a positive correlation with the BMD of the femur trabecular bone **(Figure 2F)**. A similar trend of Tregs was also observed in the BM and MLN with a significantly lower percentage of Tregs (p<0.05) in the ovx group in comparison to the sham group **(Figure 2D-E)**.

**Figure 2:**
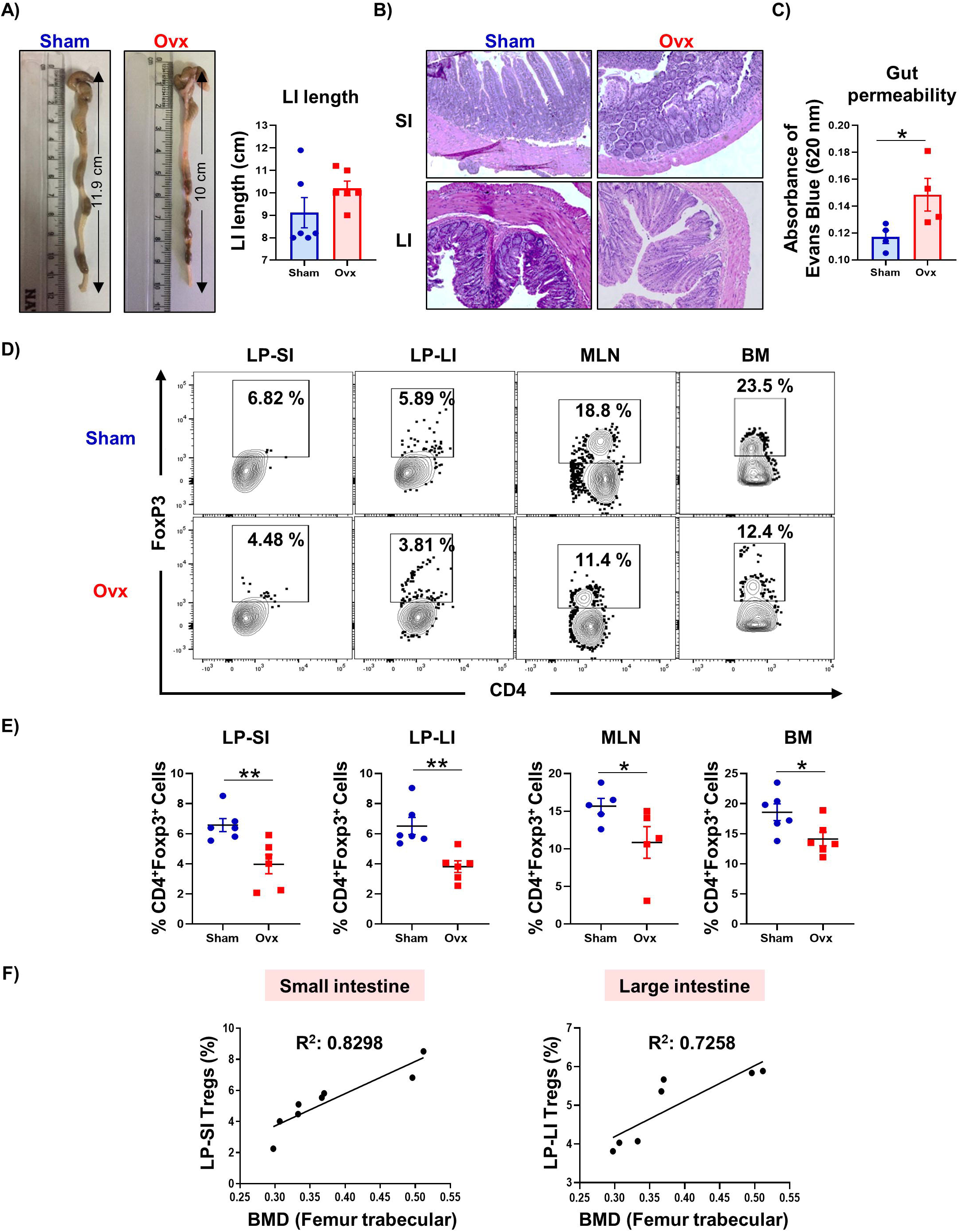
Ovariectomy results in modulation of gut-resident Treg (GTreg) cell population *in vivo*. A) Measurement of colon length in sham and ovx groups. B) Histology of small intestine (SI) and large intestine (LI) of sham and ovx group. C) Bar plot depicting absorbance of Evans Blue in the LI at 620 nm in sham and ovx group. D) Cells from the lamina propria of the small intestine (LP-SI) and large intestine (LP-LI), mesenteric lymph nodes (MLN), and bone marrow (BM) of sham and ovx groups were harvested and analyzed by flow cytometry for the percentage of Tregs. D) Dot plot depicting percentages of Tregs (CD4^+^FoxP3^+^) in LP-SI, LP-LI, MLN, and BM of sham and ovx. E) Bar graph representing percentages of Tregs in sham and ovx. F) Correlation graphs depicting the correlation of Tregs from the SI and LI with the bone mineral density (BMD). The results were evaluated using the Student t-test for paired or non-paired data, as appropriate. Values are expressed as mean ± SEM (n=6) and similar results were obtained in two independent experiments. Statistical significance was defined as *p ≤ 0.05, **p < 0.01 ***p ≤ 0.001 concerning the indicated mice group.

### 3.2 GTregs suppress osteoclastogenesis and enhance osteoblastogenesis

To investigate the effect of GTregs on bone health *in vitro*, naïve T cells were magnetically selected from the MLN (a gut-associated lymphoid tissue) and cultured under Tregs polarizing conditions **(Figure 3A)**. Following the generation of GTregs, their ability to influence the differentiation of bone cells viz., osteoclast and osteoblast was evaluated. For osteoclastogenesis, BMCs were stimulated in the presence or absence of GTregs at different cell ratios using an osteoclastogenic medium supplemented with MCSF (30 ng/ml) and RANKL (60 ng/ml). Interestingly, we observed that GTregs significantly inhibited osteoclastogenesis in a cell ratio-dependent manner, indicated by the number of TRAP-positive cells and cells with more than 3 nuclei (**Figure 3B)**. Moreover, a substantial (more than 30-fold) decrease in the area of osteoclasts in the treated groups as compared to the control was observed **(Figure 3B)**. For osteoblastogenesis, BMCs were cocultured with GTregs (BMC: GTregs; 10:1, 5:1, and 1:1) in the presence of osteogenic induction media consisting of β-glycerophosphate (10 mM) and ascorbic acid (50 µg/ml). On day 7, cells were fixed and processed for ALP staining. Remarkably, GTregs significantly enhanced osteoblastogenesis in a cell ratio-dependent manner as evidenced by the increased percentage of the stained area of ALP in the treated groups **(Figure 3C)**.

**Figure 3:**
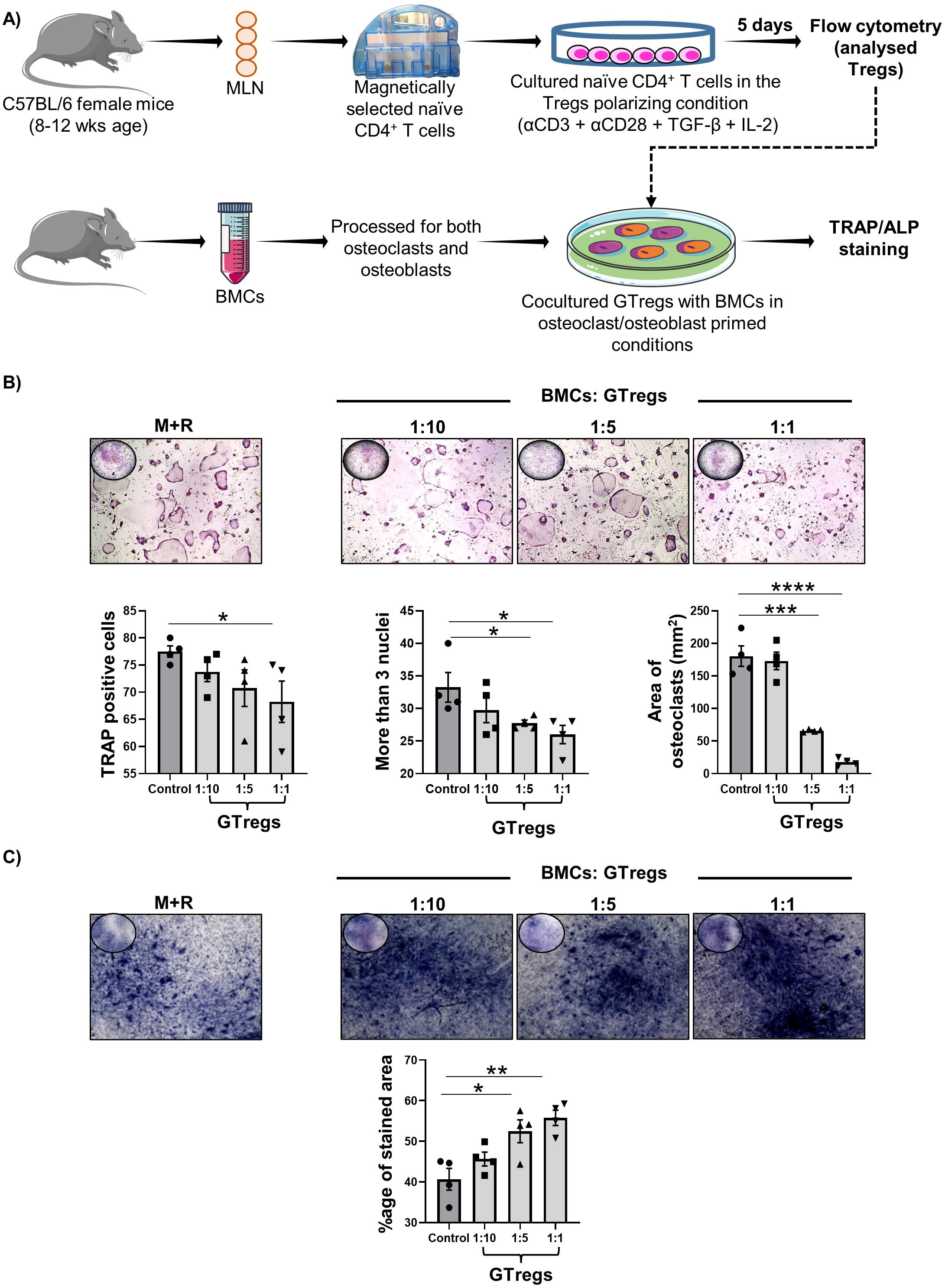
Gut-resident Tregs (GTregs) suppress osteoclastogenesis and promote osteoblastogenesis dose-dependently. A) Naïve T cells were magnetically selected from the mesenteric lymph nodes (MLN) and cultured in the Tregs polarizing conditions. After incubation for 5 days, GTregs were cocultured with the bone marrow cells (BMCs) in the presence of M-CSF (30ng/ml) and RANKL (60 ng/ml) for 4 days. Giant multinucleated cells were stained with TRAP, and cells with ≥ 3 nuclei were considered mature osteoclasts. B) Photomicrographs at 10X magnification were taken. Osteoclast differentiation was analyzed by plotting for TRAP-positive cells, the number of TRAP-positive cells with more than 3 nuclei, and the area of osteoclasts. C). Osteoblast differentiation was induced in BMCs with osteoblast induction media (OIM) with or without GTregs at different ratios for 7 days. Osteoblastogenesis was determined by alkaline phosphatase (ALP) staining and the percentage of the stained area was calculated with the help of ImageJ software. Similar results were obtained in at least two independent experiments (n≥2). Statistical significance was considered as *p≤0.05,**p≤0.01, ***p≤0.001)with respect to indicated groups.

### 3.3 Gut-resident pTregs and tTregs are dysregulated during PMO

Next, to comprehend the role distinct Treg populations play in the regulation of bone health, we further analyzed the population of both tTregs and pTregs within the Tregs population **(Supplementary Figure 1)** in various gut tissues (LP-SI, LP-LI, and MLN) and BM of both sham and ovx mice. Interestingly, we observed that ovx mice had a significantly decreased percentage of pTregs (CD4^+^Foxp3^+^NRP-1^−^) and an increased percentage of tTregs (CD4^+^Foxp3^+^NRP-1^+^) in comparison to control mice in the gut tissues i.e., LP-SI (p<0.05), LP-LI (p<0.05) and MLN (p<0.05) and BM (p<0.05) **(Figure 4A-C)**. Additionally, the proportion of pTregs in the SI and LI was positively correlated with the BMD of femur trabecular bone, while the frequencies of tTregs showed a negative correlation **(Figure 4D)**. Collectively, our results demonstrate that gut pTregs and tTregs are dysregulated during PMO and thus could play a critical role in regulating bone health.

**Figure 4:**
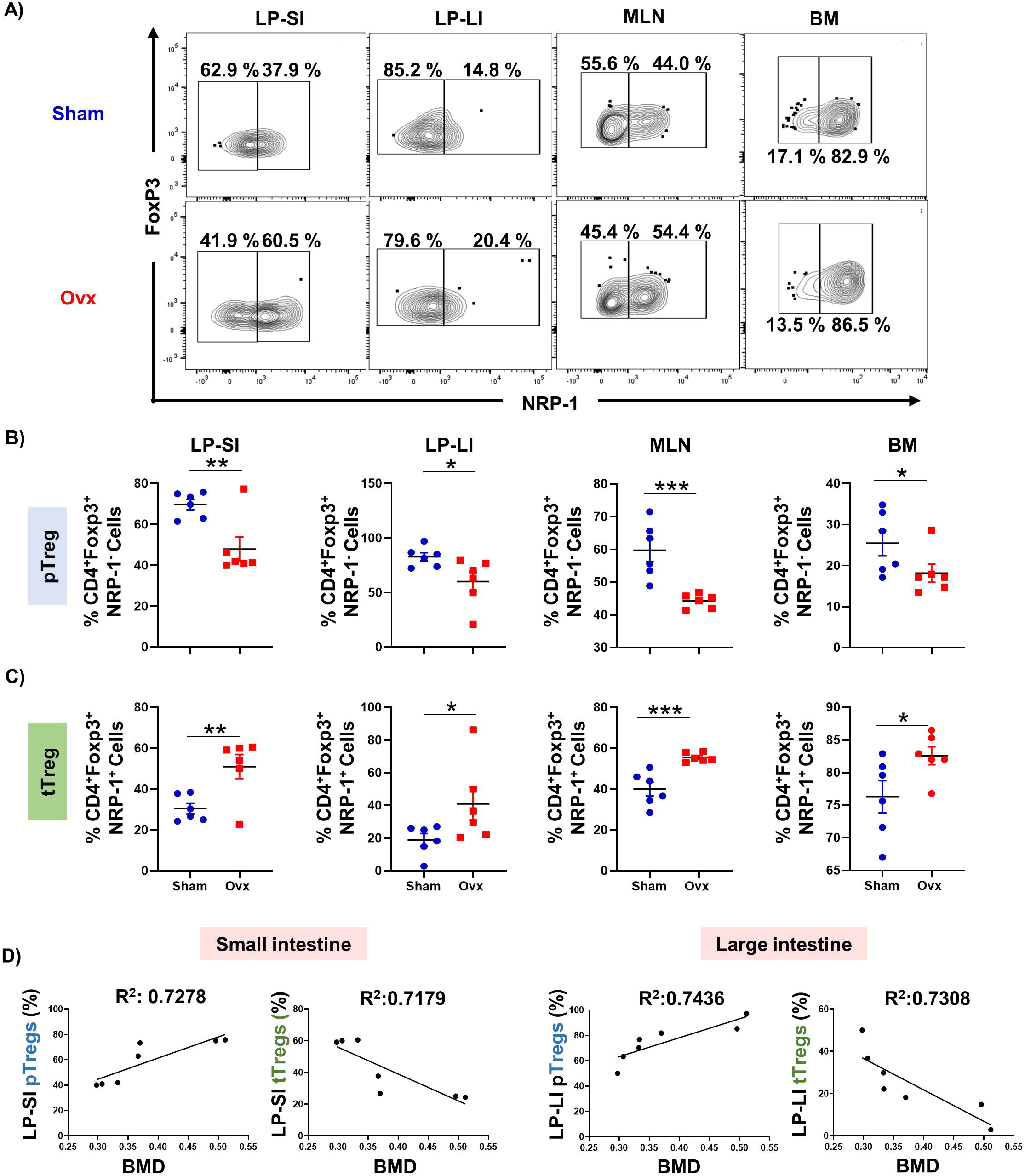
Ovariectomy alters the population of gut-resident Treg (GTreg) cell compartments. Cells from the lamina propria of the small intestine (LP-SI) and large intestine (LP-LI), mesenteric lymph nodes (MLN), and bone marrow (BM) of sham and ovx groups were harvested and analyzed by flow cytometry for the percentage of pTregs (CD4^+^Foxp3^+^NRP-1^−^) and tTregs (CD4^+^Foxp3^+^NRP-1^+^). A) Dot plot depicting percentages of pTregs and tTregs in LP-SI, LP-LI, MLN, and BM of sham and ovx. B) Bar graph representing percentages of pTregs in sham and ovx. C) Bar graph representing percentages of tTregs in sham and ovx. D). Correlation graphs depicting the correlation of pTregs and tTregs from the SI and LI with the bone mineral density (BMD). The results were evaluated using the Student t-test for paired or non-paired data, as appropriate. Values are expressed as mean ± SEM (n=6) and similar results were obtained in two independent experiments. Statistical significance was defined as *p ≤ 0.05, **p < 0.01 ***p ≤ 0.001 for the indicated mice group.

### 3.4 Probiotics administration restores the homeostasis balance of GTregs in ovx mice

Our group has reported that probiotics LR and BL alleviate bone loss ^10,26^ however, in our previous studies we haven’t studied the involvement of GTregs in probiotics-mediated amelioration of bone loss in ovx mice. Thus to validate this, we randomly divided adult female C57BL/6J mice into four groups: sham, ovx, ovx + LR, and ovx + BL. Ovx + LR and ovx + BL groups were orally administered with either LR (10^9^ CFU) or BL (10^9^ CFU) daily for 45 days **(Supplementary Figure 2).** At the end of the treatment, mice were sacrificed and various organs were harvested for analyzing the osteoimmune parameters. μ-CT and BMD data of LV-5 trabecular bones validated that LR and BL treatment to the ovx mice has significantly prevented the deterioration of the bone microarchitecture **(Supplementary Figure 2)** and also has been reported in detail in our earlier publications ^10,26^

We next evaluated the potential of these probiotics in the modulation of the GTregs. To accomplish the same, immune cells were harvested from the gut tissues (LP-SI, LP-LI, and MLN) and BM. As expected, the imbalance of the Tregs in the ovx mice had been restored by the administration of the probiotics LR and BL. LR treatment significantly enhanced the percentage of the total Tregs in the LP-LI (p<0.05), LP-SI (p<0.05), and BM (p<0.01) in comparison to the ovx mice **(Figure 5A-B)**. Similarly, BL treatment significantly enhanced the total Tregs in the LP-SI (p<0.01) and LP-LI (p<0.05) as compared to the ovx mice. However, no significant change was observed in MLNs.

**Figure 5:**
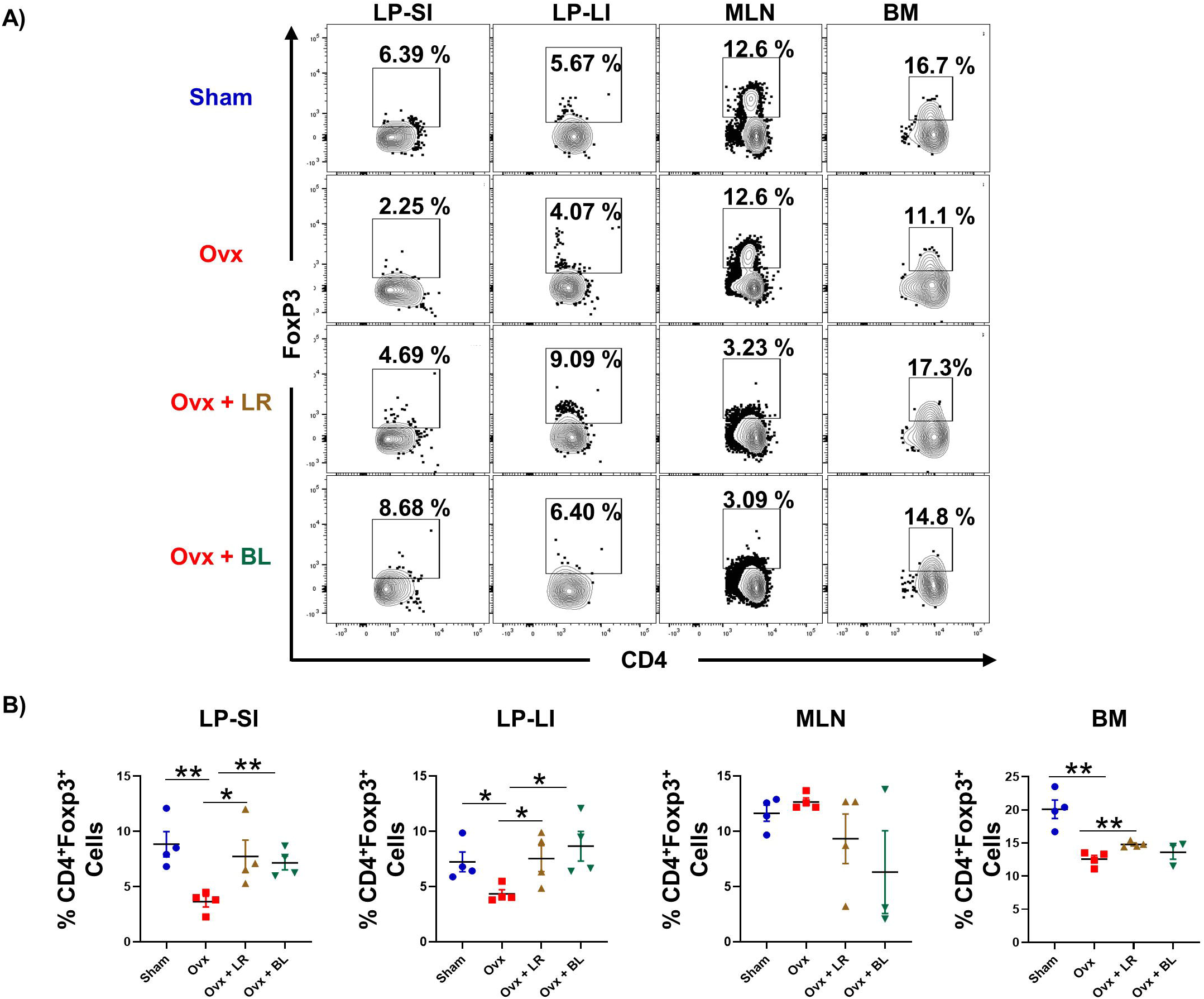
LR and BL administration restores the homeostasis balance of GTregs in ovx mice. Cells from the lamina propria of the small intestine (LP-SI) and large intestine (LP-LI), mesenteric lymph nodes (MLN), and bone marrow (BM) of sham and ovx groups were harvested and analyzed by flow cytometry for the percentage of Tregs (A) Dot plots representing the percentages of CD4^+^FoxP3^+^ Tregs in LP-SI, LP-LI, MLN and BM in sham, ovx, ovx + LR and ovx + BL groups (B) Bar graphs representing percentages of CD4+Foxp3+ Tregs in LP-SI, LP-LI, MLN, and BM. Data are reported as mean ± SEM (n=5). Similar results were obtained in two independent experiments. The statistical significance of each parameter was assessed by ANOVA followed by paired or unpaired group comparison. *p ≤ 0.05, **p < 0.01 ***p ≤ 0.001 for the indicated mice group.

Further, we examined the role of probiotics in maintaining GTreg homeostasis in the intestine of the ovx mice. We observed that LR administration significantly enhanced the population of the pTregs in the gut tissues i.e., LP-SI (p<0.05), LP-LI (p<0.01), and MLN (p<0.01) and BM (p<0.05) as compared to the ovx mice. LR treatment further decreased the proportion of the tTregs in the LP-SI (p<0.001), LP-LI (p<0.01), MLN (p<0.01), and BM (p<0.05) in comparison to the ovx mice **(Figure 6A-C)**. Similar results were observed with probiotic BL. BL administration to the ovx mice significantly increased the frequency of the pTregs in the LP-SI (p<0.05), LI-LP (p<0.05), and BM (p<0.05) along with decreasing the tTregs in LP-SI, LP-LI (p<0.05) and BM (p<0.05), as compared to the ovx mice **(Figure 6A-C)**. However, no significant change was observed in MLNs with BL administration. Collectively our data showed that the administration of probiotics LR and BL preferentially enhances gut-resident pTregs rather than tTregs thereby maintaining GTreg homeostasis in the intestine.

**Figure 6:**
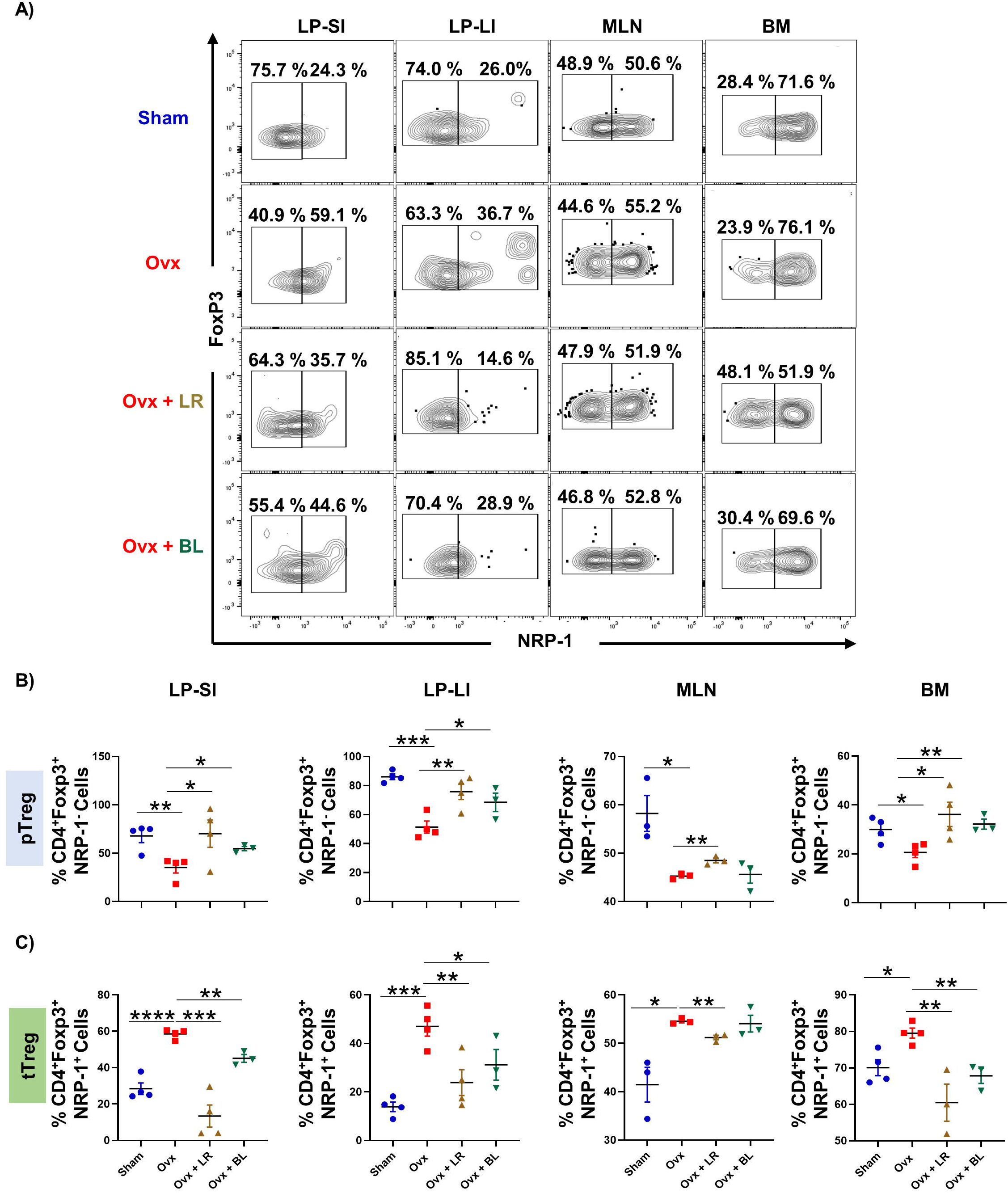
LR and BL administration modulates gut-resident pTreg and tTreg cells *in vivo*. Cells from the lamina propria of the small intestine (LP-SI) and large intestine (LP-LI), mesenteric lymph nodes (MLN), and bone marrow (BM) of sham and ovx groups were harvested and analyzed by flow cytometry for the percentage of pTregs and tTregs. A) Dot plots representing the percentages of CD4^+^FoxP3^+^NRP-1^−^ pTregs, and CD4^+^FoxP3^+^NRP-1^+^ tTregs in LP-SI, LP-LI, MLN, and BM in sham, ovx, ovx + LR and ovx + BL groups. B) Bar graphs representing percentages of pTregs in LP-SI, LP-LI, MLN, and BM. C) Bar graphs representing percentages of pTregs in LP-SI, LP-LI, MLN, and BM. Data are reported as mean ± SEM (n=5). Similar results were obtained in two independent experiments. The statistical significance of each parameter was assessed by ANOVA followed by paired or unpaired group comparison. *p ≤ 0.05, **p < 0.01 ***p ≤ 0.001 for the indicated mice group.

### 3.5 Short-chain fatty acids (SCFAs) promote differentiation of GTregs

Since probiotics are known to have their immunomodulatory effects vis SCFAs, thus we next performed targeted HPLC for the analysis of SCFAs (acetate, propionate, and butyrate) in the fecal samples collected from all the groups of mice **(Figure 7A)**. HPLC data demonstrated that ovx mice have significantly decreased amounts of SCFAs in the fecal samples compared to the controls **(Figure 7B)**. On the other hand, administration with the probiotic LR significantly enhanced the level of the acetate (p<0.05) and butyrate (p<0.05) in comparison to the ovx mice. Similarly, BL treatment significantly increased the level of all the SCFAs, acetate (p<0.01), propionate (p<0.05), and butyrate (p<0.001) as compared to the ovx mice **(Figure 7B)**.

**Figure 7:**
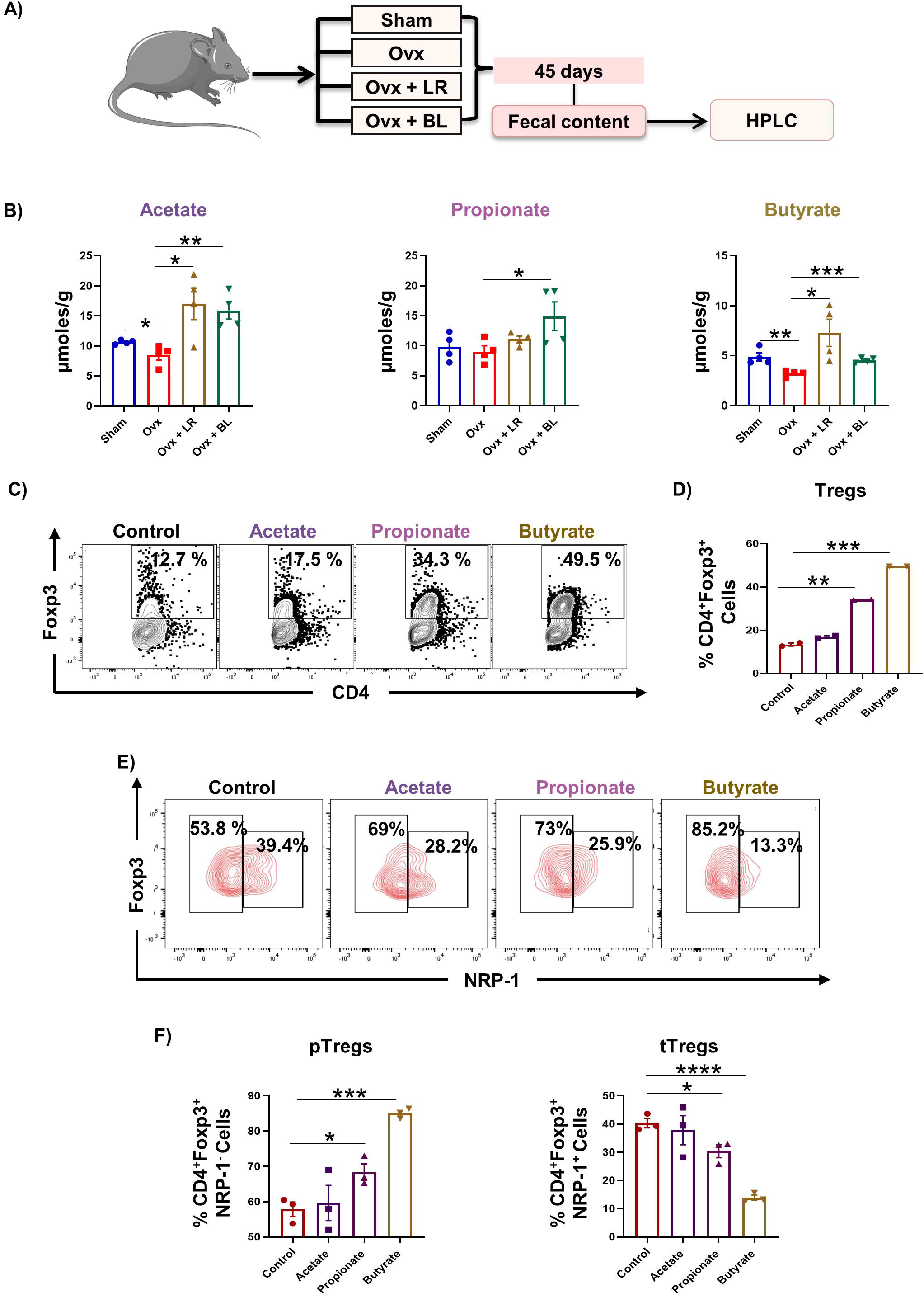
Short-chain fatty acids (SCFAs) promote differentiation of GTregs. A) At the end of 45 days, fecal samples are collected from all the groups of mice and analyzed for short-chain fatty acids (SCFAs) with the help of high-performance liquid chromatography (HPLC). B) Bar graphs representing the concentration of acetate, propionate, and butyrate in fecal samples of sham, ovx, ovx + LR, and ovx + BL groups. C) Naïve T cells were magnetic selected from the mesenteric lymph node (MLN) and cultured in the Tregs polarizing conditions in the presence of acetate, propionate, and butyrate at 0.3 mM concentration. D) Bar graphs representing the percentages of Tregs within the *in vitro* induced cells. E) Dot plots and F) Bar graphs representing the pTregs and tTregs within the *in vitro* induced Tregs. Data are reported as mean ± SEM. The statistical significance of each parameter was assessed by ANOVA followed by paired or unpaired group comparison. *p ≤ 0.05, **p < 0.01 ***p ≤ 0.001 for the indicated mice group.

Next, we estimated the role of SCFAs on the differentiation of GTregs. For the same, naïve T cells were isolated from MLNs (gut-associated/resident naïve T cells) and cultured under Tregs polarizing conditions in the presence and absence of the SCFAs (acetate, propionate, and butyrate). On day 5, Tregs were evaluated using flow cytometry. The results indicated that the SCFAs propionate (p<0.01) and butyrate (p<0.001) significantly enhanced the differentiation of Tregs **(Figure 7C-D)**. Further, we evaluated the pTregs and tTregs populations within the total Tregs. The results indicated that treatment with SCFAs propionate (p<0.05) and butyrate (p<0.01) significantly enhanced the differentiation of the pTregs along with inhibiting the differentiation of tTregs as compared to the control with butyrate being most potent **(Figure 7E-F)**. These results thereby clearly highlight that SCFAs (mainly butyrate) induce the differentiation of GTregs primarily by promoting the differentiation of pTregs.

### 3.6 SCFAs primed GTregs have enhanced anti-osteoclastogenic potential

Next, we assessed whether SCFAs have the potential to further enhance the anti-osteoclastogenic and pro-osteoblastogenic potential of GTregs. For the same SCFAs (acetate, propionate, and butyrate) primed GTregs were cocultured with BMCs at different cell ratios (10:1, 5:1, and 1:1) for both osteoclastogenesis and osteoblastogenesis. TRAP results evidenced that SCFAs-primed GTregs were more potent in significantly inhibiting osteoclastogenesis in comparison to control/unprimed GTregs **(Figure 8A-D)**. On the other hand, our ALP assay demonstrated that, in contrast, to control GTregs, SCFA-primed GTregs were unable to enhance osteoblastogenesis **(Data not shown)**. Conclusively, our above data suggest that SCFAs primed GTregs have enhanced anti-osteoclastogenic potential than control GTregs and thus could be more effective in preventing bone resorption. We also investigated the direct effect of SCFAs on the osteoclasts and observed that SCFAs significantly decreased osteoclastogenesis as indicated by the decrease in the number of TRAP-positive cells with more than 3 nuclei and area of osteoclasts **(Figure 8E-G)**. Thus SCFAs can suppress bone resorption both directly via inhibiting osteoclastogenesis and indirectly via enhancing the anti-osteoclastogenic potential of GTregs.

**Figure 8:**
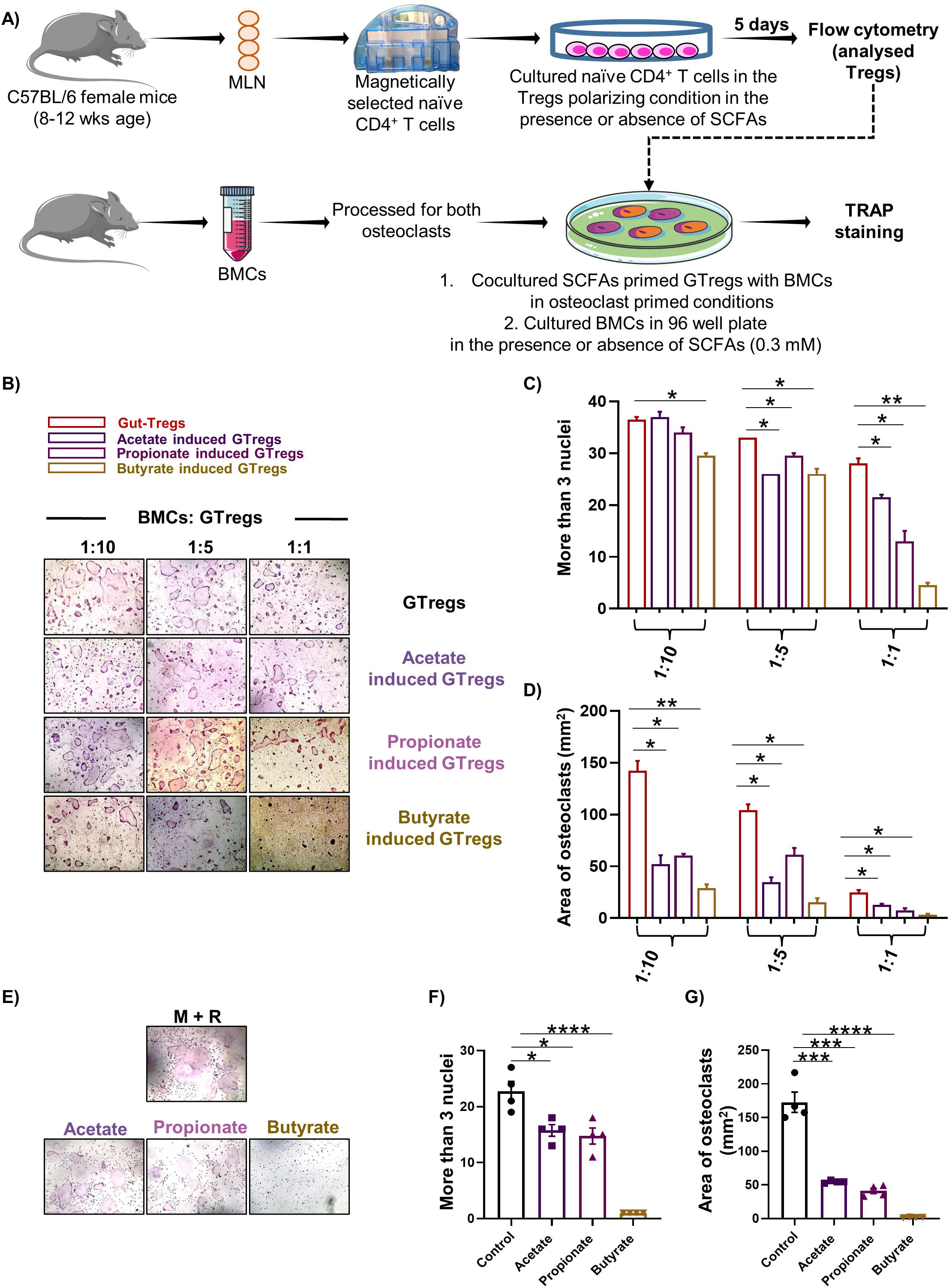
SCFAs-induced GTregs inhibit osteoclastogenesis. A). Naïve T cells were magnetically selected from the mesenteric lymph node (MLN) and cultured in the presence or absence of acetate, propionate, and butyrate at 0.3 mM concentration. (1).SCFAs-induced GTregs were then cocultured with the bone marrow cells (BMCs) in the presence of M-CSF (30ng/ml) and RANKL (60 ng/ml) for 4 days. (2) BMCs were also cultured without GTregs in the presence of MCSF and RANKL with or without SCFAs (0.3mM). B) Pictographs representing the osteoclast differentiation in the presence and absence of acetate, propionate, and butyrate-primed GTregs. Osteoclast differentiation was analyzed by plotting the TRAP-positive cells with more than C) 3 nuclei and D) area of osteoclasts. E) BMCs were induced into osteoclasts in the presence of MCSF and RANK with or without SCFAs (0.3 mM) for 4 days. Osteoclast differentiation was analyzed by plotting the TRAP-positive cells with more than F) 3 nuclei and G) area of osteoclasts. Similar results were obtained in at least two independent experiments (n≥2). Statistical significance was considered as *p≤0.05,**p≤0.01, ***p≤0.001 with respect to indicated groups.

### 3.7 GTreg’s homeostasis is perturbed in osteoporotic patients

Next, we further evaluated our results under clinical conditions. Due to the scarcity of colon tissue from osteoporotic patients, we recruited IBD patients with or without osteoporosis for colon tissue, a routine for clinical biopsy. The IBD patients were divided into three groups based on their T score calculated from the DEXA scan: Control IBD patients (without osteopenia and osteoporosis), IBD patients with osteopenia, and IBD patients with osteoporosis. Next, we isolated immune cells from the colon biopsy and analyzed them for Tregs **(Figure 9A and Supplementary Figure 3).** Interestingly it was observed that compared to the control IBD patients, the population and absolute count of Tregs were decreased in the IBD patients with osteopenia (p<0.05 and p<0.01), with a further decrease in IBD patients with osteoporosis (p<0.05) **(Figure 9B)**. Likewise to mice in humans also it was observed that the frequency and absolute counts of pTregs were significantly decreased in the IBD patients with osteopenia (p<0.05) and osteoporosis (p<0.05) compared to the IBD controls **(Figure 9B)**. Remarkably the decrease in the pTregs population is associated with the increase in the tTregs population in the IBD patients with osteopenia (p<0.05) and osteoporosis (p<0.05) compared to the control IBD patients **(Figure 9B)**. Furthermore, we observed that the frequency of Tregs in the IBD patients positively correlated with the T score of the lumbar spine **(Figure 9C)**. Similar to the mice data pTregs and tTregs showed positive and negative correlation respectively with the T score of the lumbar spine **(Figure 9C)**. Altogether our human data further corroborates our preclinical data, that osteoporosis is associated with dysregulated frequency of GTregs (decreased pTregs and enhanced tTregs).

**Figure 9:**
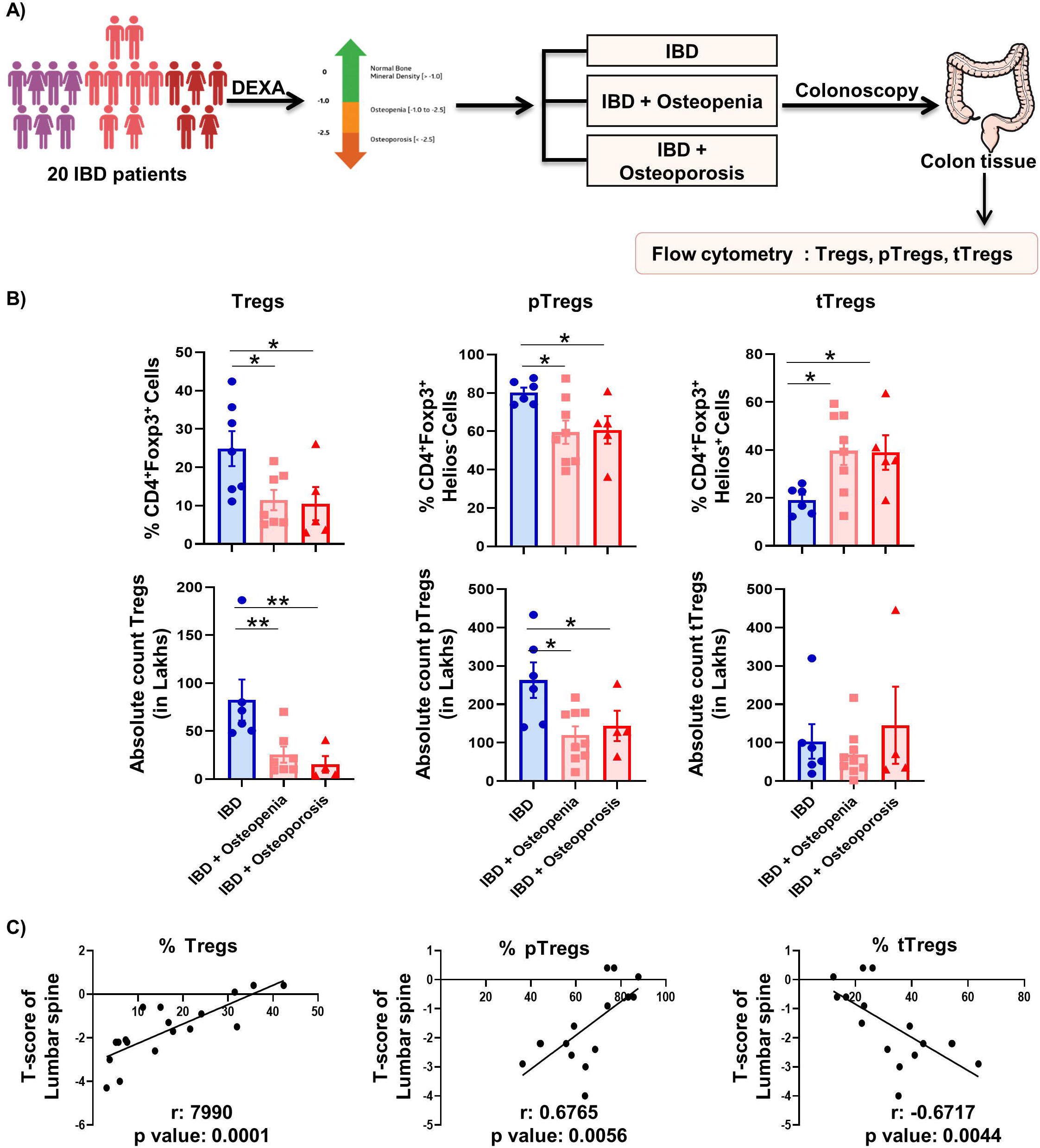
GTreg’s homeostasis is perturbed in osteoporotic patients. A) Methodology adopted for analysis of Tregs in the human colon. B) Bar graphs representing the percentage and absolute counts of Tregs (CD3^+^CD4^+^Foxp3^+^), pTregs (CD3^+^CD4^+^Foxp3^+^Helios^−^), and tTregs (CD3^+^CD4^+^Foxp3^+^Helios^+^) in the human colons. C) Correlation graphs depicting the correlation of Tregs, pTregs, and tTregs from the bone mineral density (BMD) of the Lumbar spine. The statistical significance was considered as *p≤0.05,**p≤0.01, ***p≤0.001 concerning indicated groups.

## 4.0 Discussion

Impairment in the normal bone remodeling process results in various skeletal manifestations such as osteoporosis. According to the conventional perspective, mechanical factors, endocrine problems, and metabolic disorders are the main causes of osteoporosis ^29^. However, new developments in the field of bone biology have unconditionally demonstrated the importance of the immune system in modulating inflammatory bone loss under osteoporotic conditions, a field now called “Immunoporosis” ^2–5^. Tregs are critical for bone health, especially in regulating osteoclast formation. Although the intestine is a primary site for Tregs, the role of GTregs in the pathophysiology of PMO has not been studied. Previous studies have shown that gut permeability increases in osteoporosis ^22,23^. We too observed that PMO causes histological damage to the SI and LI along with increasing gut permeability. We further observed that compared to the sham, there was a significant reduction in the frequency of Tregs in the LP-SI and LI-LI in the ovx mice. Moreover, the percentage of the Tregs in both the LP-SI and LP-LI showed a positive correlation with the BMD. This data therefore evidenced that intestinal inflammation is a manifestation of PMO, which in turn disturbs the balance of Treg cells in the intestine.

Further, we assessed the role of GTregs on bone health via regulating the activity of osteoclasts and osteoblasts. We observed that GTregs enhance osteoblastogenesis along with inhibiting osteoclastogenesis, thereby increasing bone formation. This suggests that GTregs could play a crucial role in regulating bone remodeling in PMO.

Intestinal Tregs are further divided into two subsets: pTregs and tTregs. Some earlier studies have documented the role of dysbiosis in osteoporosis ^21,30^. Since gut microbiota and their metabolites induce the differentiation of pTregs, we postulated that dysbiosis under osteoporotic conditions might modulate the balance of pTregs and tTregs in the intestine. pTregs contribute to the control of autoimmune diabetes and are highly efficient in controlling the onset of autoimmune gastritis ^31–33^. However, the role of pTregs in PMO is still unexplored. Remarkably, our data showed that, in comparison to the sham, the frequency of pTregs is significantly reduced and that of tTregs is significantly increased in the gut tissues and BM in the ovx group, supporting our hypothesis that PMO perturbs the balance of gut-resident pTregs and tTregs. Notably, a positive correlation between the BMDs (LV-5) and the frequency of pTregs along with a negative correlation with the percentage of tTregs in the gut tissues was observed.

Moving ahead, we next analyzed whether probiotics exhibit the potential to modulate the homeostatic balance of pTregs and tTregs in ovx mice. Remarkably, both the probiotics (LR & BL) enhanced the frequency of the total Tregs in the gut tissues and BM of the ovx mice. Strikingly, probiotics restored the dysregulated balance of pTregs and tTregs in the intestine of ovx mice. Bacterial metabolites such as SCFAs have immunomodulatory properties and have a key role in the development of Tregs ^34–37^. Given that both LR and BL are known to stimulate the production of SCFAs ^38–40^, we analyzed SCFA levels in the fecal samples from all groups. We found that ovx mice had lower SCFA levels, and probiotic treatment effectively restored these levels. SCFAs were further observed to promote the development of GTregs from naïve T cells. Moreover, SCFAs were shown to promote the differentiation of pTregs and decrease that of tTregs within the total Tregs population. Markedly, SCFA-primed GTregs were more effective at inhibiting osteoclastogenesis compared to SCFA-unprimed or control GTregs, leading to enhanced bone health.

Moving ahead we further corroborated our findings under clinical settings in osteoporotic subjects. Due to paucity of obtaining colon samples from normal PMO subjects, we enrolled IBD patients with osteopenia/osteoporosis, as colon biopsies of these patients are routinely available for colonoscopic/clinical evaluations. We categorized the IBD patients into three groups viz. : Control IBD patients (without osteopenia and osteoporosis), IBD patients with osteopenia, and IBD patients with osteoporosis. We analyzed Treg cell populations in these groups and observed that GTregs were significantly decreased in the colon of IBD patients with osteopenia and osteoporosis compared to the control IBD patients without osteoporosis. GTreg homeostasis was further perturbed with an increase in the frequency of the tTregs and a concurrent decrease in the frequency of pTregs in the IBD patients with osteopenia and osteoporosis compared to the IBD controls, thereby further validating our pre-clinical results in ovx mice.

Altogether, our current findings elucidate mechanisms governing the homeostatic regulation of GTregs with clinical implications. Our results indicated that gut pTregs are decreased during PMO and thus could be targeted as a feasible treatment modality for the management of PMO. Moreover, gut pTregs can be easily induced by a variety of stimuli, including microbial metabolites therefore, identifying specific metabolites and bacterial species that promote the development of gut pTregs could be instrumental in developing novel treatment strategies for the management and treatment of PMO. Summarily, our study suggests a better understanding of targeting selected population of GTregs as a novel immunotherapy in the management and treatment of inflammatory bone loss in PMO.

## Acknowledgments

AB, LS, and RKS acknowledge the Department of Biotechnology, AIIMS, New Delhi-India for providing infrastructural facilities. AB and LS thank ICMR for the research fellowship. A graphical abstract is created with the help of Biorender.

## Author Contributions

RKS contributed to the conceptualization, formal analysis, funding acquisition, investigation, project administration, resources, supervision, validation, and writing-original draft, review, and editing. AB and LS: contributed to data curation, formal analysis, methodology, and writing-original draft. DM and VA: contributed to data curation, PKM: methodology, and formal analysis. All authors reviewed the manuscript.

