## Supplementary figures and images for "Homeostatic balance of Gut-resident Tregs (GTregs) plays a pivotal role in maintaining bone health under post-menopausal osteoporotic conditions"

### Supplementary Figure S1

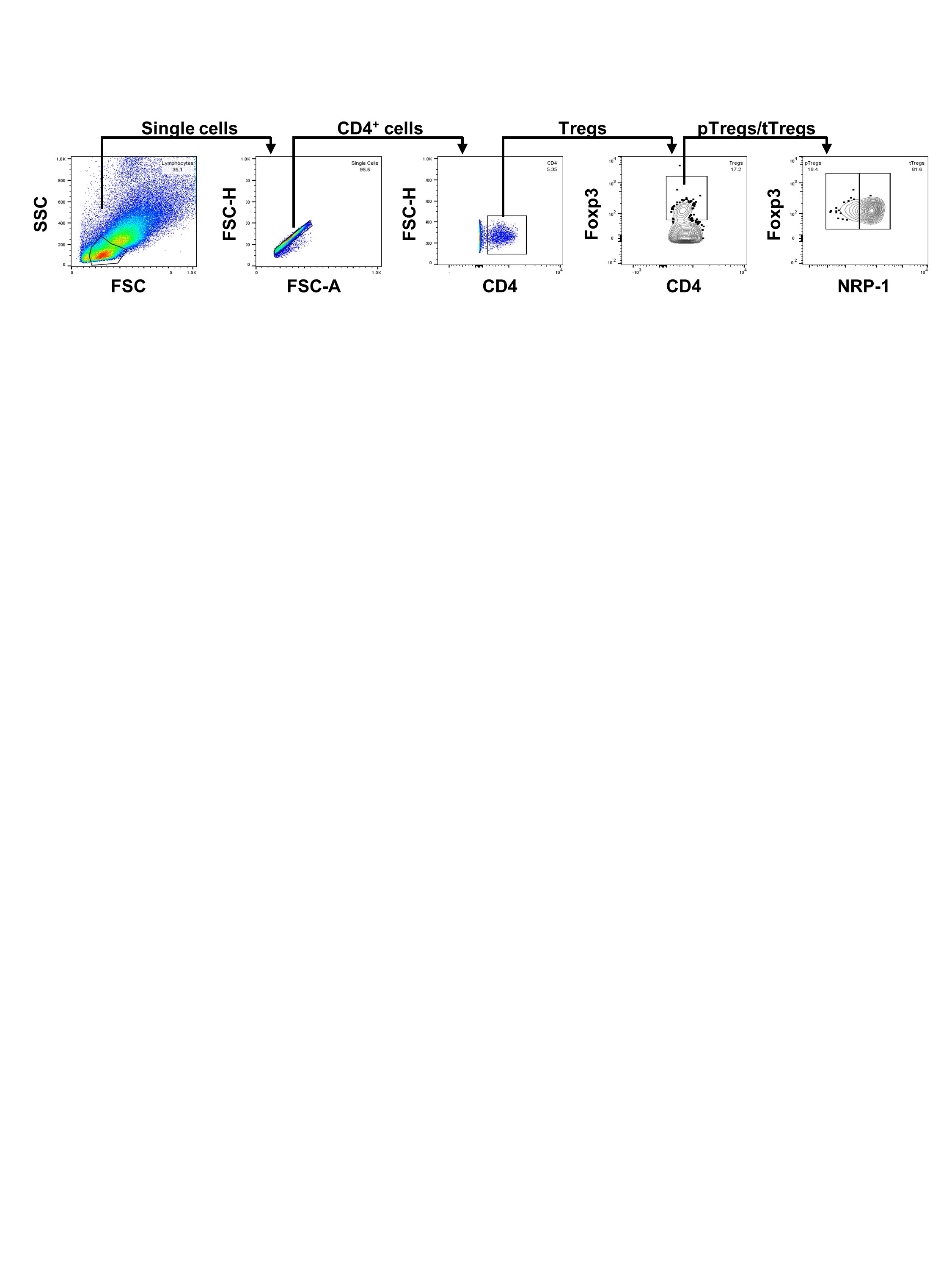

### Supplementary Figure S2

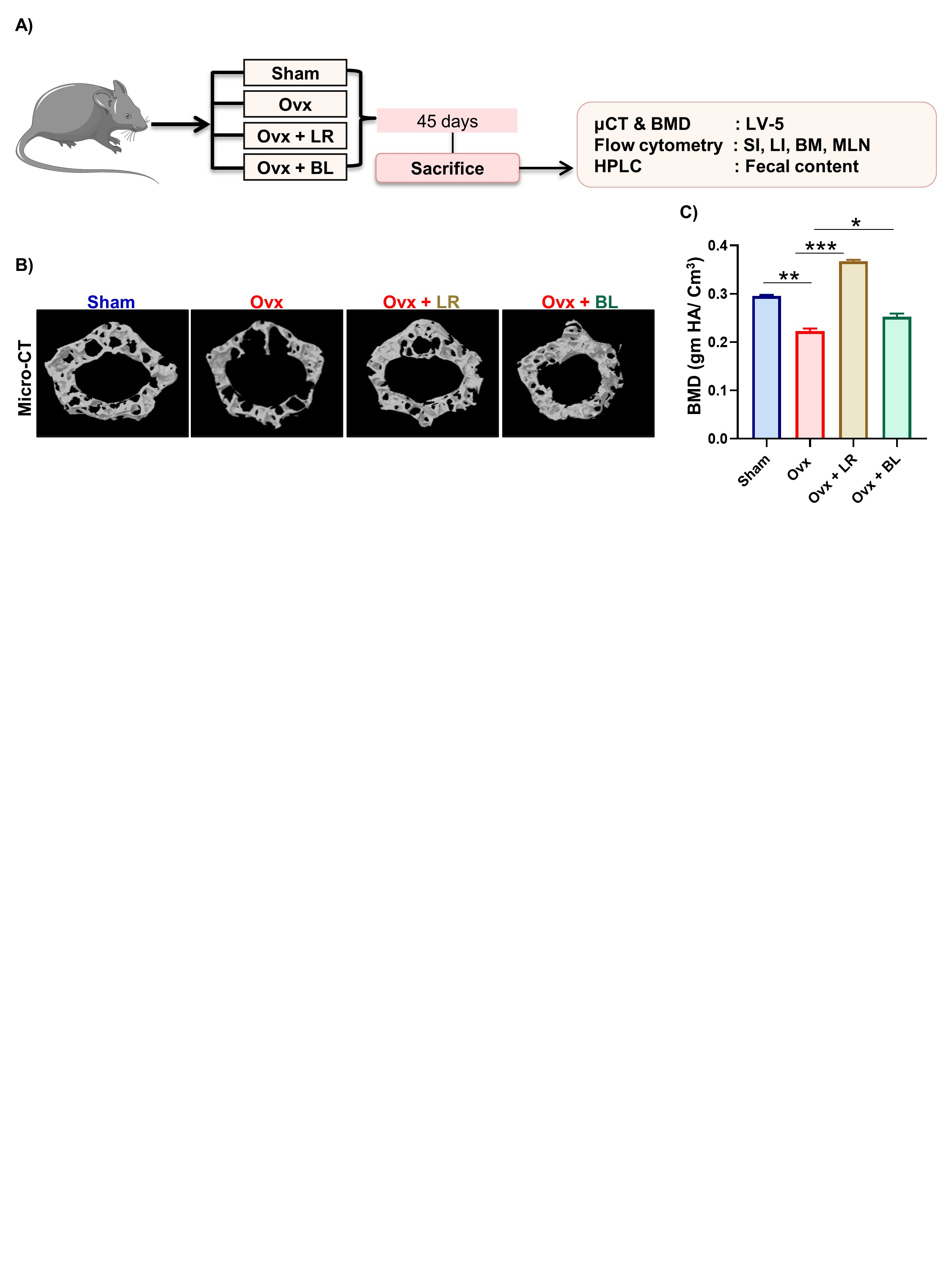

### Supplementary Figure S3

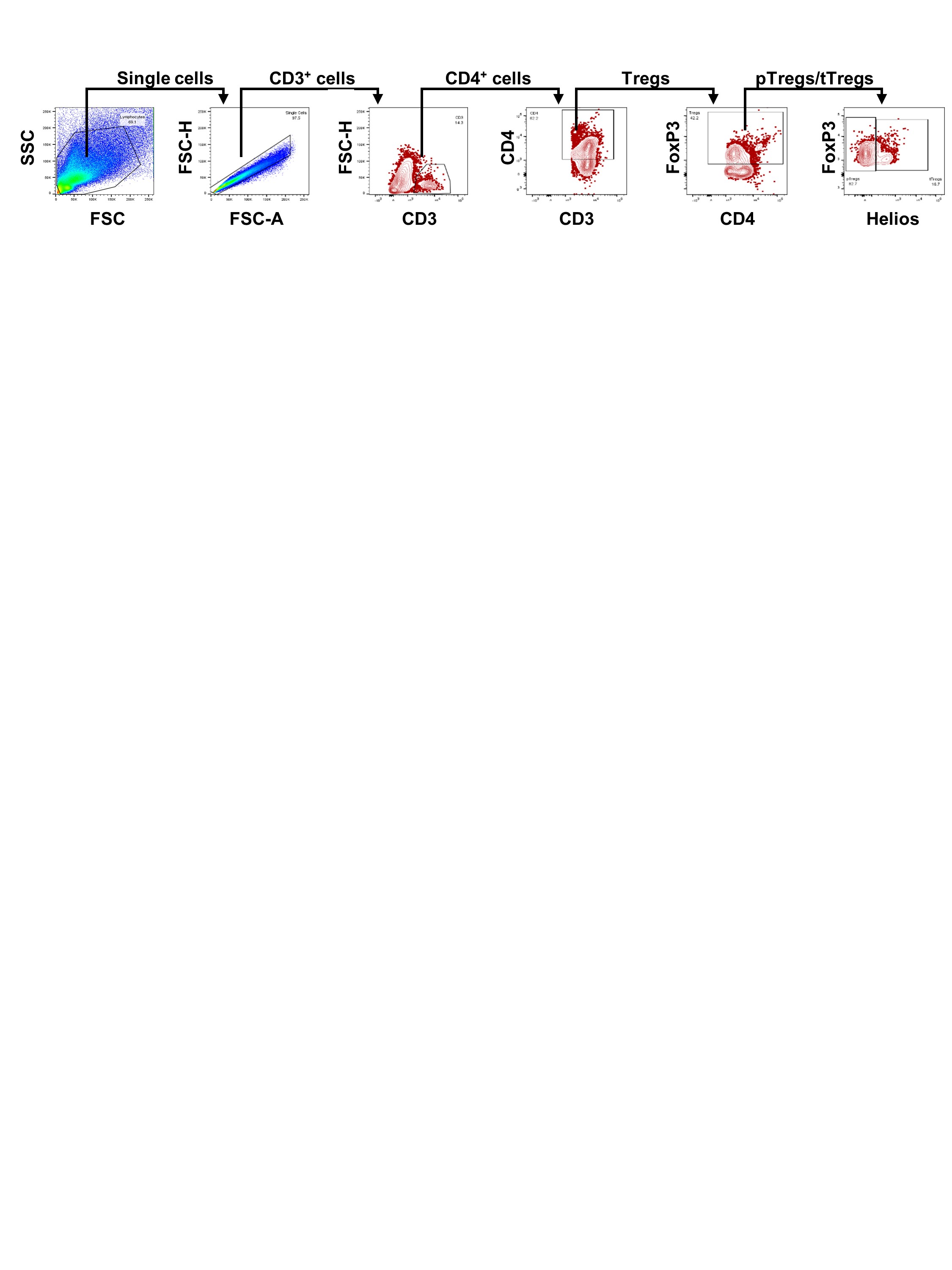
